# Epigenome-wide study uncovers tau pathology-driven changes of chromatin organization in the aging human brain

**DOI:** 10.1101/273789

**Authors:** Hans-Ulrich Klein, Cristin McCabe, Elizabeta Gjoneska, Sarah E. Sullivan, Belinda J. Kaskow, Anna Tang, Robert V. Smith, Jishu Xu, Andreas R. Pfenning, Bradley E. Bernstein, Alexander Meissner, Julie A. Schneider, Sara Mostafavi, Li-Huei Tsai, Tracy L. Young-Pearse, David A. Bennett, Philip L. De Jager

**Affiliations:** Center for Translational & Computational Neuroimmunology, Department of Neurology, Columbia University Medical Center, New York, New York, USA; Broad Institute, Cambridge, Massachusetts, USA; Picower Institute for Learning and Memory, Massachusetts Institute of Technology, Cambridge, Massachusetts, USA; Department of Neurology, Ann Romney Center for Neurologic Diseases, Brigham and Women’s Hospital and Harvard Medical School, Boston, Massachusetts, USA; Computational Biology Department, Carnegie Mellon University, Pittsburgh, Pennsylvania, USA; Department of Pathology, Massachusetts General Hospital, Boston, Massachusetts, USA; Department of Stem Cell and Regenerative Biology, Harvard University, Cambridge, Massachusetts, USA; Rush Alzheimer’s Disease Center, Rush University Medical Center, Chicago, Illinois, USA; Department of Medical Genetics, University of British Columbia, Vancouver, British Columbia, Canada; Department of Statistics, University of British Columbia, Vancouver, British Columbia, Canada; Centre for Molecular Medicine and Therapeutics, Vancouver, British Columbia, Canada

**Keywords:** Alzheimer’s disease, Tauopathy, Tau protein, Chromatin remodeling, Histone modifications, Nuclear lamina

## Abstract

Accumulation of tau and amyloid-β are two pathologic hallmarks of Alzheimer’s disease (AD). Here, we conducted an epigenome-wide association study using the H3K9 acetylation (H3K9Ac) mark in 669 aged human prefrontal cortices: in contrast to amyloid-β, tau protein burden had a broad effect on the epigenome, affecting 5,590 out of 26,384 H3K9Ac domains. Tau-related alterations aggregated in large genomic segments reflecting spatial chromatin organization, and the magnitude of these effects correlated with the segment’s nuclear lamina association. We confirmed the functional relevance of these chromatin changes by demonstrating (1) consistent transcriptional changes in three independent datasets and (2) similar findings in two AD mouse models. Finally, we found that tau overexpression in iPSC-derived neurons disrupted chromatin organization and that these effects could be blocked by a small molecule predicted to reverse the tau effect. Thus, we report large-scale tau-driven chromatin rearrangements in the aging human brain that may be reversible with HSP90 inhibitors.

## Introduction

Alzheimer’s disease (AD) is a chronic neurodegenerative disease characterized pathologically by the accumulation of amyloid-β plaques and tau tangles which leads to neuronal cell death, cognitive impairment, and, ultimately, a diagnosis of dementia. Although loci harboring genetic risk factors have been identified in genome-wide studies^1^, a large portion of late onset AD dementia risk remains unexplained, indicating the need for complementary approaches such as exploring the aging brain’s epigenome. Epigenomic alterations can be caused by genetic and non-genetic risk factors such as life experiences and environmental exposures but can also occur as a consequence of AD pathologies^2^. Hence, studying the AD epigenome may also be helpful for understanding molecular events resulting from the toxicity of AD pathologies. Evidence for epigenomic perturbations in AD was found in smaller human studies^3^, but, so far, AD-related epigenome-wide association studies of the human cortex have been limited to DNA methylation^4,5^. These studies were important in demonstrating reproducible DNA methylation changes in AD subjects, but did not distinguish explicitly between amyloid-β- and tau-related alterations. Notably, tau pathology has recently been associated with epigenetic changes in model systems. In Drosophila, tau overexpression has been shown to relax heterochromatin^6,7^. On the other hand, there is also evidence for a physiological function of nuclear tau in maintaining and regulating heterochromatin which may be lost by pathologically phosphorylated tau^8^. However, whether these mechanisms cause major chromatin alterations in the human AD brain, translate to transcriptional alterations, and are restricted to heterochromatic regions remains unknown.

Here, we studied the acetylation of the ninth lysine of histone 3 (H3K9Ac), which marks transcriptionally active open chromatin, genome-wide in the dorsolateral prefrontal cortex (DLPFC) of 669 subjects. Our data support the hypothesis that tau but not amyloid-β causes major chromatin remodeling. We mapped the location of these alterations genome-wide, characterized the architecture of these large areas of coordinated chromatin remodeling, replicated them in transcriptomic data, demonstrated that tau is sufficient to cause such chromatin rearrangement prior to tangle formation, and identified a compound that may attenuate this chromatin perturbation.

## Results

### Tau but not amyloid-β pathology has a broad effect on histone acetylation in the human brain

We studied the acetylation of histone 3 lysine 9 (H3K9Ac) in the DLPFC of 669 participants enrolled in either the Religious Order Study (ROS) or the Rush Memory and Aging Project (MAP), two longitudinal studies of aging and dementia^9,10^. Participants were not demented upon study entry. At autopsy, neuropathologic examination was performed and quantitative measurements of the density of phosphorylated tau tangles and the burden of amyloid-β were obtained (Fig. 1a). A wide spectrum of tau tangles and amyloid-β burdens with a moderate correlation of ρ=0.48 between these two AD pathologies was observed in our subjects (Fig. 1b). ChIP-seq was performed in isolated gray matter from frozen DLPFC samples to generate genome-wide H3K9Ac profiles. A median of 55 million 36 bp single-end reads were sequenced per sample (Supplementary Excel File 1). Demographic characteristics of the subjects are given in Table 1.

**Table 1:** Summary of the ROS/MAP data set. H3K9Ac and transcription data was available for 452 samples, H3K9Ac and DNA methylation data for 645 samples, and transcription and DNA methylation data for 496. All three data types were available for 449 samples.

|  | **H3K9Ac**<br>**(n=669; ChIP-seq)** | **Transcription**<br>**(n=500; RNA-seq)** | **DNA methylation**<br>**(n=729; BeadChip)** |
| --- | --- | --- | --- |
| Age (years) | 88.3 (±6.5) | 88.5 (±6.6) | 88.0 (±6.6) |
| Male gender | 233 (35%) | 189 (38%) | 267 (37%) |
| Tau protein load (%<sup>-1/2</sup>) | 2.2 (±1.4) | 2.1 (±1.3) | 2.1 (±1.4) |
| Amyloid-β load (%<sup>-1/2</sup>) | 1.5 (±1.1) | 1.5 (±1.1) | 1.5 (±1.1) |
| Pathologic AD diagnosis | 412 (62%) | 290 (58%) | 441 (60%) |
| Post mortem interval (hours) | 7.5 (±5.9) | 7.1 (±4.9) | 7.4 (±5.8) |

**Figure 1:**
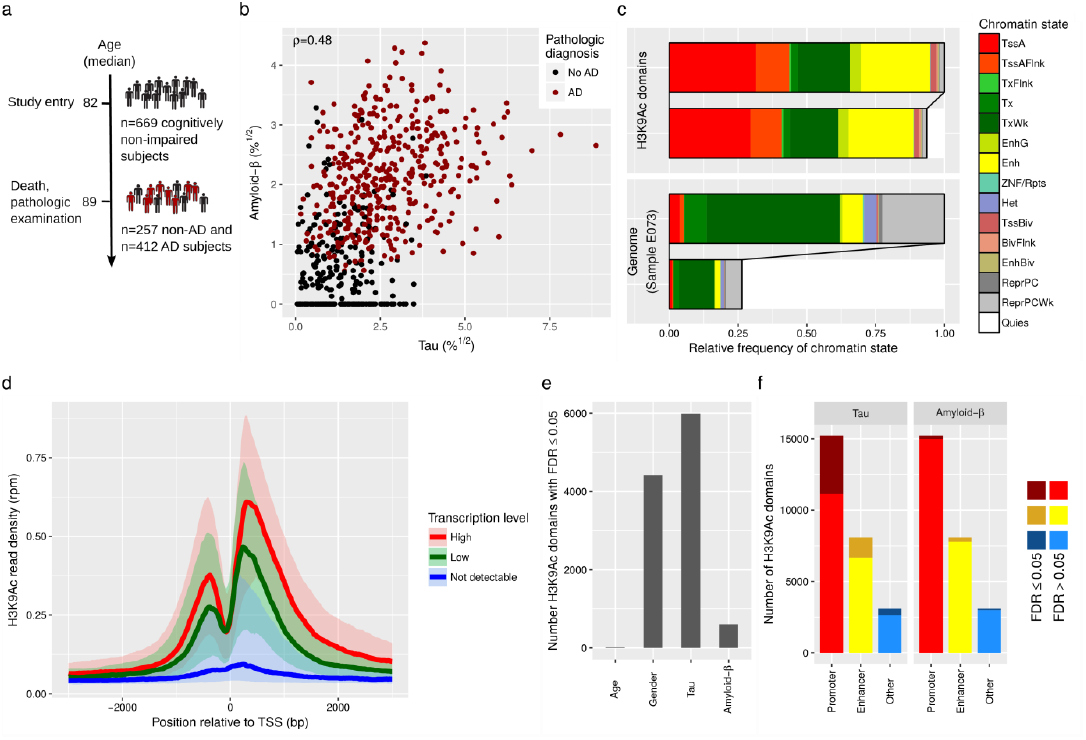
The active histone mark H3K9Ac is broadly associated with tau pathology in the human cortex. **a**, H3K9Ac was studied in the prefrontal cortex of 669 subjects. **b**, Distributions of tau and amyloid-β levels in cohort are shown. Amyloid-β peptide levels were below the detection threshold for 103 samples. **c**, Horizontal bars depict the relative frequencies of each chromatin state within the H3K9Ac domains (97 Mbp) and the whole genome (3.1 Gbp). Chromatin states were obtained from a DLPFC sample with minimal neuropathology (E073) included in the Roadmap Epigenomics Project. For better clarity, the respective upper bars depict the relative frequencies without the quiescent state. **d**, The median H3K9Ac ChIP-seq read coverage is shown on the y-axis for highly transcribed, lowly transcribed and not detectable RefSeq transcripts within ±3 kb around the transcription start site. Transcripts were grouped by median transcription values (high: x > 22 rpkm (0.95 quantile), low: 1 rpkm < × < 1.5 rpkm (0.6 - 0.65 quantiles), not detectable: x = 0). The median of the H3K9Ac read density across all samples was calculated for each transcript. Then, the median of all transcripts within each of the three groups was calculated and plotted as solid line. The 25% and 75% quantiles for each group are depicted by the transparent bands. **e**, Bars depict the number of H3K9Ac domains that were significantly associated at an FDR of 0.05 with age, gender, tau levels and amyloid-β levels. **f**, Bars depict the number of H3K9Ac domains stratified by promoter, enhancer and other domains. Dark shading indicates the number of domains whose H3K9Ac levels were significantly associated with tau or amyloid-β.

We found 26,384 H3K9Ac peaks or “domains” which showed a distinct signal above the genome-wide background in at least 100 of the 669 subjects. As with DNA methylation^4^, there is limited inter-individual variability: the average correlation of domains between two individuals was 0.98, despite the vast differences in life experiences between these older individuals. Nearly half (41%) of the genomic regions covered by H3K9Ac domains in our data were annotated as containing an active transcription start site, and more than a quarter (27%) were annotated as being in an enhancer site (Fig. 1c) based on the chromatin state annotation generated for this cortical region by the Roadmap Epigenomics Project (sample E073; a MAP subject with minimal neuropathology)^11^. We thus binned H3K9Ac peaks into “promoter” (n=15,225), “enhancer” (n=8,071) and “other” (n=3,088) domains (Supplementary Fig. 1a-d) for subsequent analyses. Using RNA-seq data from the same region in a subset of subjects (n=500), we verified the expected positive correlation between H3K9Ac and transcriptional activity (Fig. 1d).

To distinguish between tau- and amyloid-β-related epigenomic changes, we modeled both pathologies simultaneously as explanatory variables in a regression model with the H3K9Ac levels as the outcome. At a false discovery rate of 0.05, 23% of H3K9Ac domains showed an association with tau whereas only 2% were associated with amyloid-β (Fig. 1e, Supplementary Excel File 2). Only a few domains (n=88) had acetylation levels that were significantly associated with both pathologies. This is a striking and unexpected difference in the impact of tau and amyloid-β, and it may be related to the fact that tau pathology initially accumulates intracellularly and may directly affect neuronal chromatin organization, whereas amyloid-β is secreted and affects cells extracellularly. For tau, the largest proportion of associated domains was observed among promoter regions (Fig. 1f), but the estimated effect sizes were similar for domains found in promoters, enhancers and other regions (Supplementary Fig. 1e-f), indicating that the mechanism driving these associations is not specific for a type of domain. The greater average read depth at promoters probably explains the larger number of significant domains in these regions. Interestingly, known AD risk loci^1^ were not enriched in tau-associated H3K9Ac domains: out of 39 H3K9Ac domains within 50 kb of 19 known AD loci, only 9 were associated with tau, corresponding to the genome-wide relative frequency of 23% (Supplementary Table 1).

Next, we investigated whether tau-related changes in H3K9Ac led to functional consequences by analyzing our RNA-seq data: we repeated the simultaneous evaluation of amyloid and tau pathology for 18,257 out of 24,594 active transcripts that could be mapped to an H3K9Ac domain. We observed a positive but weak correlation of 0.14 between the coefficients for tau (0.12 for amyloid) from the transcription and the H3K9Ac data: in general, epigenomic and transcriptomic data showed the same direction of effect (Supplementary Fig. 1h-i) (Supplementary Excel File 3), with notable exceptions that indicate the presence of other regulatory mechanisms. The modest extent of the correlation is probably attributable to the fact that H3K9Ac is not sufficient to define a transcriptionally active region (Supplementary Fig. 1g).

### Spatial pattern of alterations in H3K9Ac indicates tau-induced remodeling of higher-order chromatin structure

Because of the large number of tau-associated H3K9Ac domains, we evaluated the spatial distribution of tau-associated domains using a “Chicago Plot” that presents the physical location, significance, and directionality of each tested domain’s association with tau (Fig. 2a, Supplementary Fig. 2a). Throughout the genome, we find large-scale genomic segments whose H3K9Ac domains were coordinately enriched for associations with tau. Unlike genome-wide SNP studies where we see very small genomic regions being associated with disease because linkage disequilibrium is observed only over ~60 kilobase pairs (kbp) in most regions of the human genome^12^, we see a clustering of disease-related epigenomic associations in certain segments of the genome that cover several megabase pairs (Mbp). These results suggest that a physical aspect of the chromosome may be implicated in disease. To define the boundaries of these large-scale segments we applied a segmentation algorithm^13^ to our data, and this method divided the genome into 178 segments within which domains display a similar tau association (Fig. 2a). The median size of these segments was 5.2 Mbp, and they contain a median of 79.5 H3K9Ac domains per segment. In contrast to tau, no similar spatial pattern was observed for domains associated with amyloid-β pathology (Supplementary Fig. 2b), suggesting a fundamental difference in the relation of these pathologies to the human cortex’ epigenome.

**Figure 2:**
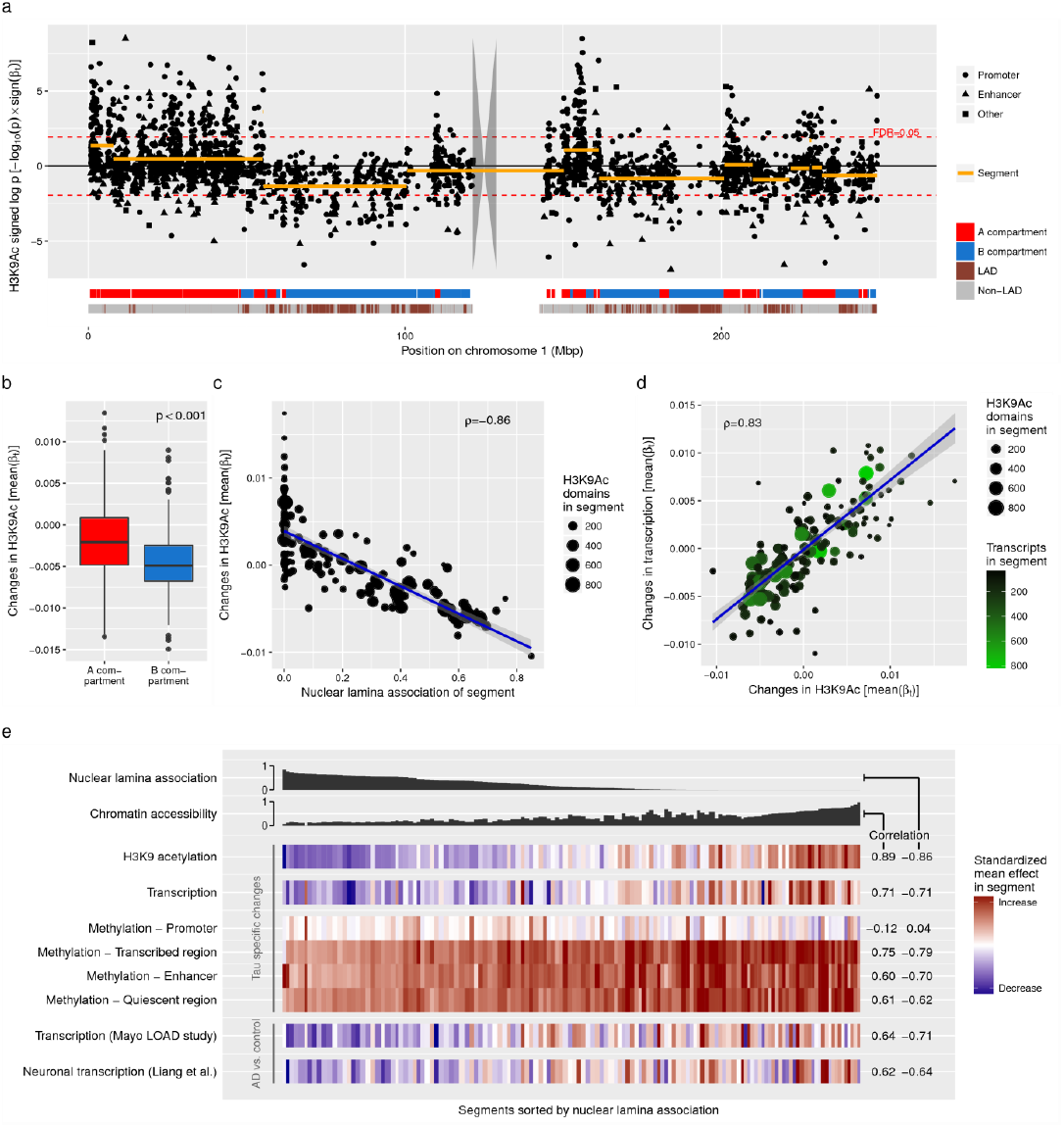
Tau-related chromatin alterations show spatial patterns. **a**, Manhattan plot depicts log-transformed p-values for the tau coefficients of all H3K9Ac domains along chromosome 1. Transformed p-values were plotted with the sign of the respective coefficient to distinguish between positive and negative associatio ns. Broad genomic segments covering H3K9Ac domains showing associations with tau similar in direction and strength are depicted as orange line segments. The gray polygon represents the chromsome’s centromere. Two ribbons at the bottom show chromatin structure related annotation. The first ribbon indicates A and B compartments. The second ribbon indicates lamina-associated domains. **b,** Boxplot shows mean tau-related changes in H3K9Ac levels observed in A compartments (n=284) and B compartments (n=223). Compartments with no or a very few H3K9Ac domains (< 5) were discarded. **c-d**, Scatter plots show the association between segments’ mean tau-related changes in H3K9Ac levels and nuclear lamina association (**c**), and mean tau-related changes in mRNA levels (**d**). Each dot represents a segment (n=138, sex chromosomes were excluded). Weighted linear regression line (blue line) and weighted Pearson correlation are depicted in the plots. Segments were weighted by the number of H3K9Ac domains. **e**, Columns of the heatmap represent segments sorted by lamina association first and then by chromatin accessibility to break ties since multiple segments were completely free of LADs. Chromatin accessibility was calculated using DLPFC specific annotation from the Roadmap Epigenomics Project (see Methods). Column widths reflect segment sizes. In rows one to six, color indicates the mean tau effect size observed in H3K9Ac, RNA transcription, DNA methylation at active TSSs, weakly transcribed regions, enhancers and quiescent regions. In the last two rows, color indicates the mean difference between AD and control samples observed in published transcription data from temporal cortex tissue (row seven) and dissected neurons (row eight).

At the megabase pair scale, chromatin is spatially organized into type A and type B compartments.^14^ Type A compartments are primarily characterized by open and type B compartments by closed chromatin. To study whether the spatial chromatin architecture underlies the observed spatial pattern of H3K9Ac alterations, we derived a compartment map from previously published Hi-C data of the cortical and subcortical plate of a human fetal brain.^15^ The genome was divided into 636 compartments and H3K9Ac domains were mapped to compartments. Figure 2a shows the locations of H3K9Ac domains and A/B compartments for chromosome 1. Genome-wide, the majority of H3K9Ac domains (70%) were located in type A compartments, but we also observed 223 type B compartments out of 507 compartments that contained at least five H3K9Ac domains. For these 507 compartments, we calculated the mean tau effect on H3K9Ac and found a significantly larger tau effect in type A compartments (Fig. 2b) indicating that the tau effect on H3K9Ac is associated with the spatial chromatin organization.

A critical element of chromatin structure is the nuclear lamina because it provides anchor points that couple chromatin to the lamina. Nuclear lamina is associated with repressive chromatin and type B compartments. Recent work in Drosophila suggests that tau can disrupt lamina function and thereby induce heterochromatin relaxation^6^. To elucidate the role of the lamina in our observations that center on euchromatin, we calculated the lamina association of our tau-defined segments as the proportion of lamina-associated domains (LADs) within a segment using previously published genome-wide DamID lamin B1 data from the human HT1080 fibrosacoma cell line^16^. Even though some LADs are cell type specific, the majority of LADs are strongly conserved, so that LADs derived from human fibroblasts are still useful for annotating whole brain tissue^16,17^. We plotted the segments’ lamina association versus the average tau effect of all H3K9Ac domains within a segment (Fig. 2c). The negative correlation (ρ=-0.86) indicates that the effect of increasing tau pathology on single H3K9Ac domains varies depending on the chromatin states of the surrounding genomic region. In fact, the 178 genomic segments within which tau associations are correlated explain 29% of the variance of the tau effects observed across all 26,384 H3K9Ac domains. Thus, these observations reflect large-scale tau-induced changes in chromatin structure.

We next evaluated our RNA-seq data and found that each segments’ mean tau effects obtained from the RNA-seq data was highly correlated (ρ=0.83) with the mean tau effects from the H3K9Ac data (Fig. 2d), reflecting a shift in average gene transcription that is related to the burden of tau pathology and is concordant with the segments’ chromatin alterations. However, in line with the weak correlation of alterations in H3K9Ac and RNA levels at the single transcript level, only 2% of the variance of tau-related effects in RNA-seq data could be explained by the segments defined with the chromatin data. This suggests that, while large-scale epigenomic changes explain some of the effect of tau on the transcriptome, other regulatory mechanisms must also be influenced by this neuropathologic process.

We then speculated that other epigenetic marks may show similar tau-related patterns: in evaluating our DNA methylation data from the same brain region^4^, we found consistent evidence for a perturbation of chromatin architecture. Specifically, we binned CpG’s into four groups using the tissue specific reference chromatin state map from the Roadmap Epigenomics Project^11^ (sample E073): active transcription start sites, enhancers, weakly transcribed regions and quiescent regions. These four chromatin states cover most of the CpGs in our data set, and we repeated the segment-based analysis for each group of CpG’s. As shown in Fig. 2e, a negative correlation between tau effects on DNA methylation and nuclear lamina association was observed in enhancers, quiescent regions and weakly transcribed regions, which is in agreement with the paradigm that high methylation of gene bodies is a feature of active genes^18^. The variance of tau effects in DNA methylation data that could be explained by the H3K9Ac derived segments ranged between 6% for enhancer, 4% for transcribed regions, and 1% for quiescent regions. By contrast, no distinct correlation was observed at active transcription start sites (explained variance < 1%), in line with previous work reporting that promoter regions are spared from altered methylation levels in the presence of other diseases like leukemia^19^.

To validate our findings in an independent set of samples, we assessed a published transcriptomic data set from the temporal cortex of subjects from a late onset AD (LOAD) case-control study^20^. Because quantitative amyloid-β and tau burdens were not available, we compared samples with AD diagnosis (n=202) to controls (n=90; non-AD and non-progressive supranuclear palsy (PSP) diagnosis) and calculated the mean AD-related change in transcription over the genes within each of the 178 genomic segments defined using our H3K9Ac data. Although we could not distinguish the tau from the amyloid-β effects in these data, we did confirm the presence of a strong negative correlation (ρ=-0.71) between the segments’ nuclear lamina association and AD-related transcriptional changes (Fig. 2e).Similar to our RNA-seq data, 4% of the variance of AD effects in the transcriptomic data of the LOAD study could be explained by the segments. Nevertheless, as in our RNA-seq data, the mean tau-related changes in the H3K9Ac data were positively correlated (ρ=0.65) with the mean AD-related changes in the transcription data from this LOAD study, validating our observation. The genomic locations of the 178 segments, their annotation and the average tau or AD effects from the different data sets are provided in Supplementary Excel File 4.

Since the data analyzed so far were generated from bulk tissue, it is critical to assess whether the observed alterations may be driven by changing cell type compositions. We estimated the proportion of NeuN^+^cells using the DNA methylation profiles^21^ and, as expected, observed a significant association (p=0.02) between the proportion of NeuN^+^cells and pathologic AD diagnosis indicating that cell type composition might be a confounder. However, the coefficients for tau derived from a model adjusted for the proportion of NeuN^+^cells fitted on a subset of subjects that have both DNA methylation and H3K9Ac data were highly correlated (ρ=0.99) with the tau coefficients from the unadjusted model (Supplementary Fig. 5h). Similarly, neither tau coefficients nor p-values changed noticeable when adjusting for transcription levels of cell type markers as outlined in the Supplementary Notes. While we cannot exclude confounding by changing cell type compositions completely^22^, these results indicate that varying cell type compositions are probably not the main driver of our observations and we decided to further investigate which cell type(s) may be effected by tau pathology.

### Altered transcription due to tau-related chromatin remodeling occurs in neurons

*A priori*, neurons are the most likely candidate cell type since intracellular phosphorylated tau is known to accumulate and form tangles in neurons. To explore this hypothesis, we repurposed a gene transcription data set of laser-capture microdissected neurons from the superior frontal gyrus of individuals with AD (n=23) and control subjects (n=11)^23^. Since tau and amyloid-β loads were not reported in these subjects, we estimated the AD effect instead of the tau effect for each gene. The AD effects were then averaged within each of the 178 genomic segments defined in our H3K9Ac data, and the correlation with the segment’s nuclear lamina association was calculated. Again, we observed a negative correlation of ρ=-0.64 (Fig. 2e), and the correlation of these neuronal data with the average tau effect in our H3K9Ac data was ρ=0.67. Similar to the tissue transcriptomic data, 2% of the AD effect variance were explained by the segments. Interestingly, the original report of these neuronal data specifies that the investigators selected neurons for laser capture that were lacking neurofibrillary tangles; this indicates that the observed AD-related transcriptional changes are occurring early in pathogenesis, prior to the accumulation of tangles^23^. Since the transcriptional changes consistent with our epigenomic changes are found in purified neurons, they are likely to be cell-autonomous for neurons; however, we cannot rule out at this time that other cell types may also be affected.

### H3K9Ac data from tau mouse models suggest similar structural changes in the murine brain epigenome that involve the nuclear lamina

To explore whether the epigenomic changes that we found in human brain are recapitulated in mouse models known to accumulate tau, we generated hippocampal H3K9Ac ChIP-seq profiles from two different mouse models at (1) an early time point and (2) a late stage of neurodegeneration (Supplementary Table 2). Specifically, we studied 6 and 11 months old mutant tau mice (*MAPT* P301S), which start to accumulate phosphorylated tau in neurons by 6 months^24^. Wild-type mice of the same age were used as controls. The second mouse model was the CK-p25 model, which is characterized by increased amyloid-β levels early after p25 induction followed by increased tau phosphorylation and neuronal loss at later stages^25,26^. Three months old CK-p25 mice were studied 2 weeks and 6 weeks after p25 induction and were compared to CK littermate controls. In total, we found 44,165 H3K9Ac domains across the epigenome of both mouse models. Tau mice showed less significantly altered H3K9Ac domains compared to the CK-p25 mice, especially at the early stage of 6 months (Fig. 3a). Chicago plots did not show the explicit spatial pattern observed in human cortex, possibly because of the low sample size of 3 replicates per time point. We then classified the mouse H3K9Ac domains into two bins: “close to lamina” (if domain <50 kb from a LAD) or “distant to lamina” based on published DamID lamin B1 data from mouse embryonic fibroblasts^16^. As anticipated, we observed smaller differences between AD mice and control mice in H3K9Ac domains that were located in the proximity of LADs (Fig. 3b-c). This observation was consistent for both mouse models and more distinct at the later time points during which the pathology has accumulated more extensively, confirming our observations from the human cortex where the effect of tau was smaller in segments enriched with LADs.

**Figure 3:**
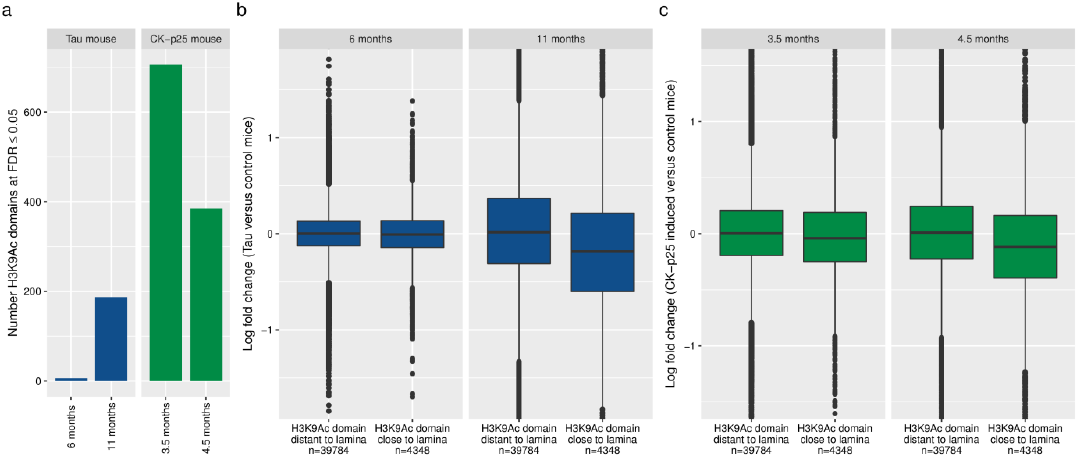
H3K9Ac alterations in AD mouse models reflect spatial pattern observed in the human cortex. **a**, Bar plot shows the number of domains with significant different H3K9Ac levels observed in the tau mouse model (blue bars) and the CK-p25 mouse model (green bars) compared to respective control mice for different time points. CK-p25 mice were 3 months old when p25 was induced. **b**,**c**, Boxplots depict differences in H3K9Ac levels between tau mice (**b**) or CK-p25 mice (**c**) and respective control mice separately for H3K9Ac domains distant to and close to LADs. The median observed log fold changes at lamina-free and lamina-associated H3K9Ac domains differed by 0.20 (p ≤ 10^−16^) at 11 months (0.01 at 6 months) for the tau model, and by 0.13 (p ≤ 10^−16^) at 4.5 months (0.04 at 3.5 months) for the CK-p25 model.

### Induced neurons overexpressing *MAPT* demonstrate chromatin structure alterations

Given the evidence from neuronal expression data that the observed changes of chromatin structure are likely to be occurring in neurons, we assessed whether overexpression of the 4R isoform of *MAPT*, which encodes the tau protein, in forebrain neurons derived from human induced pluripotent stem cells (iNs) could recapitulate the epigenomic changes that we found in our cortical H3K9Ac data. With fAD mutation, this model system induces AD-related features such as the intracellular accumulation of phosphorylated tau^27^. To characterize the epigenomic changes in this *in vitro* model system, we used the Assay for Transposase-Accessible Chromatin with sequencing (ATAC-seq) to map genome accessibility^28^. In contrast to anti-H3K9Ac ChIP-seq, ATAC-seq generates reads at all open regulatory elements where the Tn5 transposase can access exposed DNA. We therefore expect to observe ATAC-seq peaks in the vicinity of nucleosomes carrying the H3K9Ac mark that we targeted with the ChIP-seq assay, and additional peaks at enhancers and insulators that might not be marked by H3K9Ac.

All iNs were derived from the same iPSC line, and we observed 40,637 ATAC domains that have a distinct peak in at least 4 of our 18 samples. Each sample consisted of 2-3 pooled individual wells of transfected iNs; 9 samples represented independent transductions of *MAPT*-overexpressing iNs. The remaining 9 samples were control iNs transduced with a GFP overexpressing construct. Samples were generated in 3 experimental batches of independent differentiations. ATAC domains were annotated with chromatin states of hESC-derived neurons (sample E010) from the Roadmap Epigenomics Project^11^ (Fig. 4a) and clustered into four groups: transcription start site (TSS), TSS flanking region, enhancer and other domains (Supplementary Fig. 4a-c). As expected, most of the TSS domains (91%) overlapped one of the H3K9Ac domains from our whole brain tissues, but only 17% of the enhancer domains were included in our set of cortical H3K9Ac enhancer domains (Fig. 4b) indicating that H3K9Ac is not an explicit enhancer marker. Assessing these domains in the other direction, the majority of cortical H3K9Ac domains were overlapped by at least one ATAC domain (Fig. 4c). Thus, while there are important differences between the two sets of profiles, there is good overlap, particularly in TSS where the bulk of the tau effect is observed in our H3K9Ac tissue-level data.

**Figure 4:**
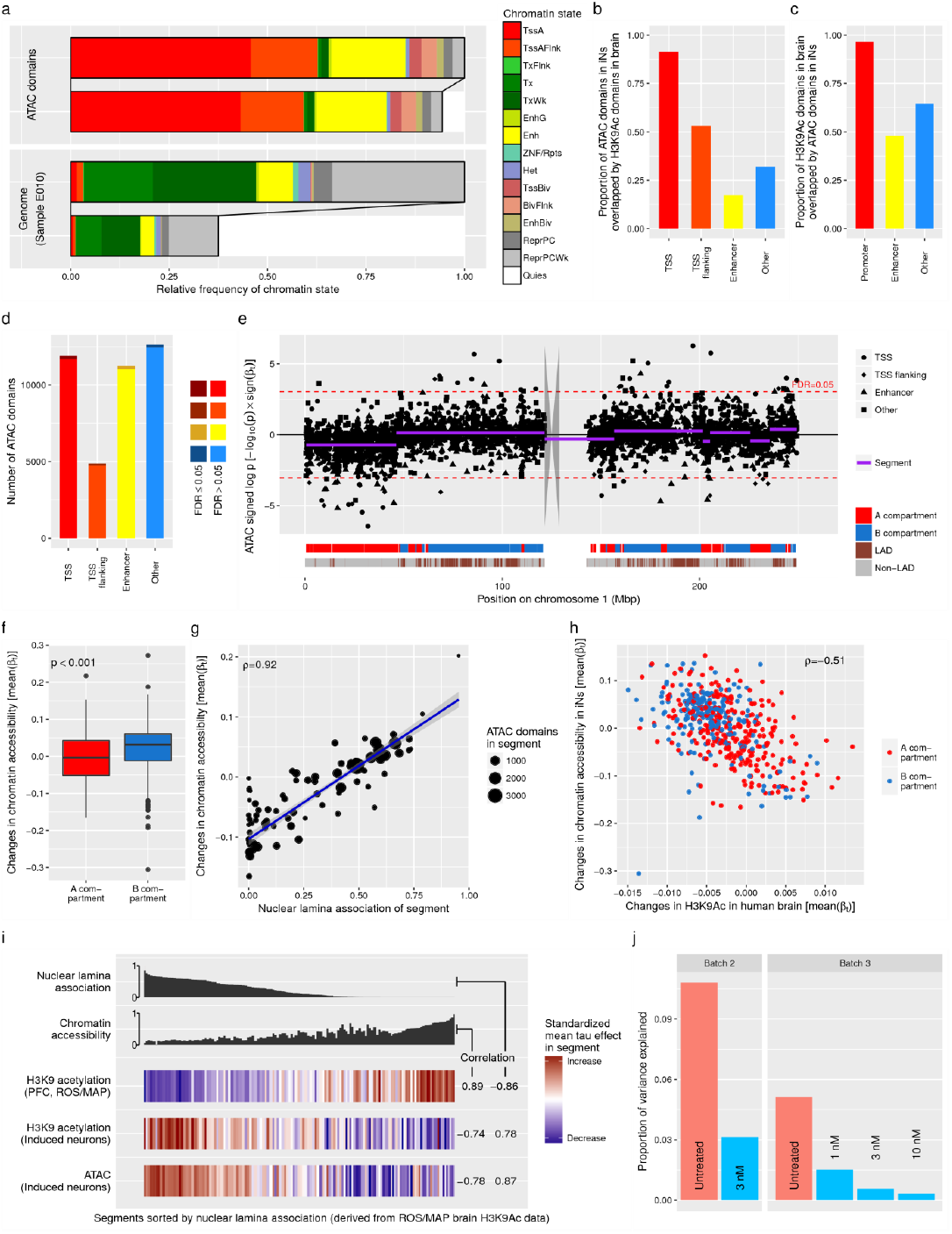
Tau induces chromatin alterations in iPSC-derived neurons. **a**, Horizontal bars depict the relative frequencies of each chromatin state within the ATAC domains (31 Mbp) and the whole genome (3.1 Gbp). Chromatin states were obtained from H9 derived cultured neurons (E010) included in the Roadmap Epigenomics Project. For better clarity, the respective upper bars depict the relative frequencies without the quiescent state. **b**, Bars depict proportion of ATAC domains in neurons that were overlapped by an H3K9Ac domain in the DLPFC. **c**, Bars depict proportion of H3K9Ac domains in the DLPFC that were overlapped by an ATAC domain in neurons. **d**, Bars depict the number of ATAC domains stratified by TSS, TSS flanking, enhancer and other domains. Dark shading indicates the number of domains whose chromatin accessibility differed significantly between *MAPT* overexpressing neurons and controls. **e**, Manhattan plot depicts the log-transformed p-values for differences between *MAPT* overexpressing neurons and controls of all ATAC domains along chromosome 1. Transformed p-values were plotted with the sign of the test statistic to distinguish between increased and decreased chromatin accessibility. Broad genomic segments covering ATAC domains showing similar changes in chromatin accessibility are depicted as purple line segments. The gray polygon represents the chromsome’s centromere. Two ribbons at the bottom show chromatin structure related annotation. The first ribbon indicates A and B compartments. The second ribbon indicates lamina-associated domains. **f,** Boxplot shows mean tau-related changes in chromatin accessibility observed in A compartments (n=306) and B compartments (n=283). Compartments with no or a very few ATAC domains (< 5) were discarded. **g**, Scatter plot shows association between segments’ nuclear lamina association and mean change in chromatin accessibility. Each dot represents a segment (n=92, sex chromosomes were excluded). Weighted linear regression line (blue line) and weighted Pearson correlation are depicted in the plots. Segments were weighted by the number of ATAC domains. **h**, Each dot represents either an A (n=284) or B (n=221) compartment derived from fetal human brain Hi-C data. For each compartment, the mean tau-related change in H3K9Ac levels in the human DLPFC on the x-axis is plotted versus the mean change in chromatin accessibility between *MAPT* overexpressing neurons and controls on the y-axis. Compartments with less than 5 H3K9Ac domains or less than 5 ATAC domains were discarded. **i**, Columns of the heat map represent the 138 genomic segments (sex chromosomes were excluded) derived from the DLPFC H3K9Ac data and were first sorted by lamina association and then by chromatin accessibility. The color in the first row indicates the mean tau effect size observed in the DLPFC H3K9Ac data as shown in Fig. 2e. The second and third rows depict the differences in H3K9Ac and chromatin accessibility observed between *MAPT* overexpressing neurons and control neurons. **j**, Bars depict the variance of differences between *MAPT* overexpressing and respective control iNs that can be explained by segments. Blue bars indicate that the *MAPT* overexpressing iNs were treated with 17-DMAG.

In total, 747 out of 40,637 ATAC domains showed a difference in their chromatin accessibility at an FDR of 0.05 when comparing the *MAPT* overexpressing iNs to control iNs infected with a non-relevant vector (Fig. 4d). The Chicago plots in Fig. 4e and Supplementary Fig. 2c depict large genomic segments of predominantly concordant tau-related changes; segments were defined using the same segmentation algorithm that we used for the cortical H3K9Ac data, and are indicated by purple segments. In iNs, we found 99 segments that explained 11% of the variance of tau effects observed for single ATAC domains (Supplementary Excel File 6). As seen in the H3K9Ac data, the tau effect differed significantly between type A and type B compartments (Fig. 4f), and the average tau effect of the segments correlated (r=0.92) with the proportion of LADs in the segment (Fig. 4g) suggesting the presence of changes in chromatin arrangement. These results were found consistently in each of the three batches (Supplementary Fig. 4d-j). Interestingly, in contrast to the H3K9Ac changes observed in human and in mouse brain tissue, segments containing many LADs showed a large positive average tau effect indicating that chromatin relaxation preferentially occurred in lamina-associated segments in this *in vitro* system. Concordantly, when looking at the human fetal brain A/B compartments, the tau effect observed in human brain H3K9Ac data was negatively correlated with the tau effect in iNs (Fig. 4h). To verify that the model system presents a tau effect that is the inverse of what we see in the human cortex, we repeated the iN experiment with anti-H3K9Ac ChIP-seq and estimated the effect size of tau for each of the 34,988 detected H3K9Ac domains. Then, using the 178 genomic segments derived from the cortical H3K9Ac data and calculated the segments’ average tau effect in (i) iN H3K9Ac data and (ii) iN ATAC data to compare it to the brain tissue H3K9Ac data (Fig. 4i, Supplementary Excel File 4). The H3K9Ac and ATAC data from iN’s consistently indicate that tau is associated with chromatin opening in genomic segments with a high fraction of lamina association. Thus, segments of the genome are significantly affected by tau pathology *in vivo* and *in vitro*, but the direction of effect is reversed in relation to tau pathology, which accumulates over decades in the human brain and over 14 days in our *MAPT* overexpressing iN model system.

### The Hsp90 inhibitor 17-DMAG is a candidate drug for attenuating tau-related alterations of chromatin structure

To find putative drugs with the potential to protect neurons from tau toxicity, we scanned the Connectivity Map data base^29^ for compounds whose gene expression signature was negatively correlated with our H3K9Ac signature for tau pathology derived from ROS/MAP cortical profiles. As described in the Methods, H3K9Ac domains were mapped to genes based on transcription start sites inside the domains. Domains without a transcription start site were discarded. Out of 1309 compounds in the database, the N-terminal Heat Shock Protein 90 (Hsp90) inhibitor 17-(dimethylaminoethylamino)-17-demethoxygeldanamycin (17-DMAG, alvespimycin) achieved the smallest p-value according to the database’s search algorithm and the fifth smallest correlation coefficient (Supplementary Excel File 7). Further, geldanamycin, another Hsp90 inhibitor is also highly significant in this analysis (p=0.0002). Hsp90 and other heat shock proteins are involved in protein folding, and Hsp90 has been previously evaluated in the context of Alzheimer’s disease^30,31^.

To assess the effect of 17-DMAG, we treated three independent cultures of *MAPT* overexpressing iNs with 17-DMAG at a concentration of 3 nM for 24 hr before harvesting (on day 31), based on published studies^32^. The differences between the treated *MAPT* overexpressing iNs and the respective controls were calculated for each of the 40,637 ATAC domains defined in iN. The variance of these tau-induced differences was then decomposed into between-segment and within-segment variance using the 99 tau-associated chromosomal segments derived for the iNs using ATAC-seq data. The prediction is that if 17-DMAG attenuates the tau effect on chromatin organization, the variance explained by the segments (i.e. the between-segment variance) should decrease for the 17-DMAG treated iNs. In our experiment, we observed such a protective effect of 17-DMAG (Fig. 4j, Supplementary Table 5). We then confirmed these results in a second experiment and a dose response curve: triplicates of *MAPT*-overexpressing iN cell cultures with 17-DMAG at concentrations of 1 nM, 3 nM or 10 nM. As controls, we cultured *MAPT* overexpressing iNs and control iNs in DMSO solution (Supplementary Table 4). While our experiments indicate that 17-DMAG may protect neurons from chromatin alterations related to tau (Fig. 4j, Supplementary Table 5), further studies with different lines of iNs are necessary to replicate the effect of 17-DMAG and to explore the interplay between 17-DMAG and tau pathology.

## Discussion

Our epigenome-wide association study revealed that tau pathology is associated with broad changes in the brain’s epigenome. We observed large genomic segments of several megabases within which H3K9Ac domains showed similar tau-associated gains or losses of histone acetylation. At the megabase pair scale, the genome is organized into two major types of compartments, A and B, defined by patterns observed in Hi-C interaction maps^14^. Type A compartments are characterized by euchromatin which is transcriptionally active whereas the inactive type B compartments exhibit high chromatin density. These two major types of compartments are essential for chromatin organization^33^ and cell identity^34^, and impose a cis- and trans-correlation structure on epigenetic marks^35^. We demonstrated that the effect of tau on H3K9Ac differed between type A and type B compartments and that the segments observed in the Chicago plot (Fig. 2a) reflected the higher-order chromatin structure. Thus, tau appears to be preferentially affecting certain elements of nuclear architecture and chromosomal organization.

The nuclear lamina is a key element in the spatial organization of chromatin and is associated with inactive B compartments. Previous work in Drosophila suggested that the lamin nucleoskeleton is disrupted in tauopathies, which causes heterochromatin relaxation and mediates neuronal cell death^6,7^. In line with this hypothesis, we observed a strong correlation between the effect of tau in a genomic segment and the nuclear lamina association of that segment. However, we measured the euchromatic mark H3K9Ac and it remains unclear how this mark relates to heterochromatin relaxation.

The association of tau pathology with alterations in chromatin architecture is not specific to the H3K9Ac chromatin mark and has functional consequences: we replicated the presence of the segments with coordinated alterations in relation to tau pathology in both DNA methylation and transcriptomic data from the same persons. Further, we find the same pattern of results in publically available transcriptomic data from a different cohort, and a second dataset from laser-captured neurons further refines the observation by implicating a specific cell population. Even though the pattern was less distinct in the various transcriptomic data sets, demonstrating the presence of these transcriptional perturbations in relation to tau is important in understanding the consequences of these alterations. Not all epigenetic changes directly translate to altered transcription. In addition, if chromatin alterations induce the transcription of epigenetically silenced genes, transcription may occur at a low level in a tissue sample, and we may thus be under-estimating the extent of transcriptional alterations in tissue-level RNA-seq data.

We employed AD mouse models and *MAPT* overexpressing iNs to further explore our findings and to address some of the limitations of our human cortex study. Looking at the results from our iN experiments, we demonstrated that *MAPT* overexpression is sufficient to induce chromatin reorganization in neurons. Further, this event occurs prior to tangle formation and neuronal cell death in our model system: perturbation of epigenomic architecture may therefore be an early event in tau pathology. This is an important point that we cannot address with our cross-sectional human H3K9Ac brain data. In the CK-p25 mouse model, we detected altered H3K9Ac levels depending on the proximity of the H3K9Ac domain to the nuclear lamina two weeks after p25 induction. For the same mouse model at the same time point, a study reported altered localization of lamin in hippocampal neurons indicating a dispersion of the nuclear lamina membrane that could affect the chromatin organization^39^. This effect was stronger after 4 weeks in line with our observations after 6 weeks and preceded apoptosis^39^. Finally, our observation in the neuronal expression data further supports the hypothesis that tau-induced epigenomic changes are an early event that occurs downstream of pathologic tau accumulation, but before neurofibrillary tangles develop since the data was generated from neurons free of tangles^23^.

Interestingly, we observed large positive tau effects on H3K9Ac in lamina free regions of the human cortex and mouse brain, but the direction of the association was reversed for iNs. This is an intriguing result and suggests a complex but definitive interaction between tau and chromatin structure. iNs grow *in vitro* in the absence of the three dimensional context of the cerebral cortex and do not interact with non-neuronal cells. The inverse directionality that we see at an early time point after *MAPT* overexpression could be an initial response of iNs to a rapid accumulation of tau (compared to accumulation over decades in the human brain), in line with recent work suggesting that nuclear tau also has a physiological function in assembling and maintaining heterochromatin^8^. Importantly, the segmental pattern and the association of the tau effect with the proportion of LADs in the segments is seen in both the human cortex and iNs, suggesting the involvement of the same mechanism. Thus, while more work is required to delineate the molecular events happening *in vivo* and *in vitro*, iNs overexpressing tau may be a useful model system with which to investigate and target tau-induced epigenomic changes in neurons. This is illustrated by our testing of an Hsp90 inhibitor that is predicted to block the effects of tau based on our brain data: 17-DMAG attenuates the chromatin alterations induced by tau in iN’s.

Overall, we have characterized changes throughout the epigenome of the human AD brain and showed that tau induced alterations of the spatial chromatin organization. The effect of these alterations propagate into the transcriptome. Our data supports the hypothesis that the nuclear lamina and the transcription factor CTCF are key elements in mediating tau toxicity *in vivo* and *in vitro*, and we demonstrated that iNs are a useful model system to further study tau-associated chromatin changes and to screen for drugs such as 17-DMAG that might prevent or reverse these chromatin alterations.

## Author Contributions

P.L.D., D.A.B., T.L.Y.-P., A.M. and B.E.B. conceived the study. C.M., E.G., A.T., S.E.S., B.J.K., A.T. and R.V.S. conducted experiments. J.A.S. and D.A.B. and contributed post mortem brain tissues. E.G. and L.-H.T. contributed mouse models. S.E.S. and T.L.Y.-P. contributed neuronal models. H.-U.K., J.X. and A.R.P. analyzed data. H.-U.K., E.G., S.M., T.L.Y.-P., D.A.B. and P.L.D. interpreted data and designed follow-up experiments. H.-U.K. and P.L.D. wrote the manuscript with contributions from all co-authors.

## Acknowledgements

This work has been supported by NIH grants U01 AG046152, R01 AG036836, R01 AG015819, R01 AG017917, R01 AG036547. L.-H. T. has been supported by NIH/NINDS/NIA (RO1 NS078839) and the Robert A. and Renee R. Belfer Family Foundation. The Mayo Clinic Alzheimer’s Disease Genetic Studies were led by Dr. Nilüfer Ertekin-Taner and Dr. Steven G. Younkin, Mayo Clinic, Jacksonville, FL using samples from the Mayo Clinic Study of Aging, the Mayo Clinic Alzheimer’s Disease Research Center, and the Mayo Clinic Brain Bank. Data collection was supported through funding by NIA grants P50 AG016574, R01 AG032990, U01 AG046139, R01 AG018023, U01 AG006576, U01 AG006786, R01 AG025711, R01 AG017216, R01 AG003949, NINDS grant R01 NS080820, CurePSP Foundation, and support from Mayo Foundation.

## Methods

### ROS/MAP cohort and pathologic characterization

All participants were enrolled in the Religious Orders Study (ROS)^9^ or the Memory and Aging Project (MAP)^10^. Studies were approved by the Institutional Review Boards of Rush University Medical Center and Partners Healthcare. Participants were not demented at the time of enrollment and agreed to donate their brain upon death. Amyloid-β and tau tangles were assessed postmortem as previously described^40,41^. Briefly, brains were cut into 1-cm-thick coronal slabs and immersion fixed in 4% paraformaldehyde. Tissue blocks from 8 brain regions (hippocampus -CA1/subiculum-, angular gyrus, entorhinal, superior frontal, dorsolateral prefrontal, inferior temporal, anterior cingulate, and calcarine cortices) were embedded in paraffin and sectioned at 20μm. Paraffin-embedded sections were immunostained for amyloid-β using 1 of 3 monoclonal anti-human antibodies: 4G8 (1:9000; Covance Labs, Madison, WI), 6F/3D (1:50; Dako North America Inc., Carpinteria, CA), and 10D5 (1:600; Elan Pharmaceuticals, San Francisco, CA). Paired helical filament (PHF) tau tangles were labeled with an antibody specific to phosphorylated tau (AT8, Thermoscientific, Waltham, MA, USA). A computerized sampling procedure combined with image analysis software was used to calculate the percentage area occupied with amyloid-β and the density of PHFtau tangles. Composite scores were computed for overall amyloid-β burden and PHFtau tangle density by averaging the scores obtained from the eight brain regions. Amyloid-β and tau tangle scores were square root transformed for better statistical properties.

### H3K9Ac ChIP-seq

We identified the Millipore anti-H3K9Ac mAb (catalog # 06-942, lot: 31636) as a robust mAb. 50mg of gray matter was dissected on ice from biopsies of the DLPFC of the ROS/MAP cohorts. The tissue was minced and crosslinked with 1% formaldehyde at room temperature for 15mins and quenched with 0.125M Glycine. The tissue was then homogenized in cell lysis buffer using the Tissue Lyser and a 5mm stainless steel bead. Then the nuclei were lysed in cell lysis buffer and chromatin was sheared by sonication. Chromatin was incubated overnight at 4C with the H3K9Ac antibody and purified with protein A sepharose beads. The final DNA was extracted and used for Illumina library construction following usual methods of end repair, adapter ligation and gel size selection. Samples were pooled and sequenced on the Illumina HiSeq (36 bp single-end reads).

### H3K9Ac ChIP-seq data pre-processing and peak detection

Single-end reads were aligned by the BWA algorithm against the human reference genome GRCh37^42^. Peaks were detected for each sample individually by MACS2 using the broad peak option and a stringent q-value cutoff of 0.001^43^. Pooled genomic DNA of seven samples was used as negative control. A combination of different ChIP-seq quality measures were employed to remove low quality samples^44^: samples that did not reach (i) ≥ 15×10^6^ unique reads, (ii) non-redundant fraction ≥ 0.3, (iii) cross-correlation ≥ 0.03, (iv) fraction of reads in peaks ≥ 0.05 and (v) ≥ 6000 peaks were removed. After quality control, 669 out of 712 samples remained (Supplementary Excel File 1). We defined our H3K9Ac domains by calculating all genomic regions that were detected as a peak in at least 100 (15%) of our 669 samples. Regions neighbored within 100 bp were merged and very small regions of less than 100 bp were removed, resulting in 26,384 H3K9Ac domains. Domains were annotated with chromatin states obtained from an unaffected DLPFC sample included in the Roadmap Epigenomics Project^11^ (E073, core 15-state model) and clustered into promoter, enhancer or other domains as described in the Supplementary Notes.

### Statistical analysis of H3K9Ac ChIP-seq data

For each sample and H3K9Ac domain, we counted the number of ChIP-seq reads that fell into the domain^45^. Read counts *Y* were assumed to follow a negative binomial distribution *Y ~ NB(μ,θ)* with mean *μ* and dispersion parameter *θ*. A log-linear regression model

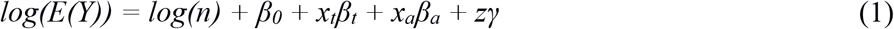

was fitted for each H3K9Ac domain separately using the R package MASS. The offset *log(n)* accounts for different total numbers of reads *n* (summed over all domains) per sample. Coefficients *β*_*t*_ and *β*_*a*_ model the effect of the observed square root-transformed tau and amyloid-β levels on the H3K9 acetylation levels. Coefficients were tested by Wald tests and the false discovery rate (FDR) was calculated to account for multiple testing. Vector *γ* models the biological covariates age and gender as well as the technical covariates post mortem interval, batch and cross-correlation, which is a measure of the efficiency of the ChIP.

### Statistical analysis of transcription and methylation data

ROS/MAP RNA-seq data from the DLPFC were generated as previously described^46^ and can be downloaded from Synapse (Synapse: syn3388564). Briefly, after sequence alignment, transcript abundances were estimated by RSEM using GENCODE v14 as reference transcriptome and reported as rpkm values (reads per kilobase of transcript per million reads)^47^. Transcripts with rpkm values larger or equal to 2 in at least 25% of the samples were considered as active in the human brain and kept in the data set (n=24,594 transcripts). A linear regression model was fitted for each transcript with the log2-transformed rpkm values as outcome variable and tau and amyloid-β as explanatory variables. The model was adjusted for age, gender, RIN score (RNA integrity number), post mortem interval, study index (ROS or MAP) and logarithmized total number of aligned reads.

DNA methylation data were generated as described elsewhere^4^ and can be downloaded from Synapse (Synapse: syn3157275). Illumina 450K bead arrays were processed by the Illumina GenomeStudio to obtain relative methylation levels (“beta values”). Relative methylation levels were assumed to follow a beta distribution and modeled by CpG-wise beta regression models using the probit function as link function^48^. Tau, amyloid-β, age, gender, post mortem interval, batch and bisulfite conversion rate were used as covariates.

Gene expression data from the Mayo LOAD study^20^ was downloaded at Synapse (Synapse: syn3157225)^49^. The neuronal expression data set was downloaded from Gene Expression Omnibus (GEO: GSE5281)^23^. Analyses of these data sets are described in the Supplementary Notes.

### Determining genomic segments of concordant tau-related changes

Tau p-values from model (1) were log-transformed and the signs of the respective tau coefficients *β*_*t*_ were assigned to the transformed p-values to account for the direction of the association. The circular binary segmentation algorithm^13^ implemented in the R package DNAcopy was then applied to the transformed p-values with the corresponding genomic positions of the H3K9Ac domains. We used the default value of *α=0.01* to accept new change-points. After the segments were determined, we calculated the weighted mean of all tau effect sizes *β* _*t*_ within the segment using the coefficients’ inverse standard deviation as weights. Mean effect sizes were obtained in the same manner for the other data sets shown in Fig. 2e and 4g by calculating the weighted average of the respective coefficients for either tau, Alzheimer’s disease diagnosis or *MAPT* overexpression. Supplementary Excel File 4 contains all 178 segments and the mean effect sizes from all data sets. Small segments (< 20 H3K9Ac domains or < 20 active transcripts) and segments on sex chromosomes were excluded when calculating correlation coefficients reducing the number of segments to 138. Genomic segments defined on the ATAC-seq data from induced neurons were identified in the same manner. All 99 ATAC-seq derived segments are given in Supplementary Excel File 6. Exclusion of small segments (< 20 ATAC domains) and segments on sex chromosomes reduced the number of segments to 89.

### A/B compartment and lamina-associated domain annotation

Human fetal brain A/B compartments were obtained from a published Hi-C data set (GEO: GSE77565) as described in the original publication.^15^ Briefly, Pearson correlation matrices were calculated from the intra-chromosomal observed/expected 40 kbp binned interaction matrices. For each chromosome, the sign of the first principal component obtained from the correlation matrix was used to categorize bins as type A or B compartment.

Genomic segments were annotated with their lamina association using published DamID data for Lamin B1 in human fibroblasts (GEO: GSE22428).^16^ Probes of the DamID array were classified into nuclear lamina bound or unbound in the original work. We re-mapped the probes against genome GRCh37 and defined the lamina association of a segment as the ratio of nuclear lamina bound probes to all probes in the segment. To sort segments in Figures 2e and 4i, chromatin accessibility was calculated from chromatin state annotation of sample E073 (DLPFC) from the Roadmap Epigenomics Project. Chromatin accessibility was defined as the fraction of the segment that was annotated with any chromatin state other than heterochromatin, repressed polycomb, weak repressed polycomb or quiescent/low. These four chromatin states exhibited the lowest DNA accessibility measured by DNase-seq^11^. As expected, chromatin accessibility was negatively correlated with lamina association in the DLPFC (ρ=-0.89): that is, segments of more open chromatin were less likely to include a LAD.

### Mouse models

All mouse work was approved by the Committee on Animal Care of the Division of Comparative Medicine at MIT and complied with the relevant ethical regulations. Adult (3 months old) female double-transgenic CK-p25 mice^25^ and their respective control littermates were used for the experiments. Brain tissue was collected at either 2 or 6 weeks after p25 induction. For the Tau (P301S) mouse model^24^, adult female 6 months old and male 11 months old transgenic mice with age- and gender-matched wild-type mice as controls were used. Upon dissection the tissue was flash-frozen in liquid nitrogen. No animals were excluded from the study. Hippocampal tissue was extracted and ChIP-seq was applied as described for the human brain tissue using the Abcam antibody ab#4441 (lot numbers GR196840-1 and GR207546-1). Supplementary Table 2 summarizes the number of replicates generated for the different time points.

### Pre-processing and statistical analysis of the mouse H3K9Ac ChIP-seq data

The murine H3K9Ac ChIP-seq data was pre-processed similar to the human DLPFC ChIP-seq data. BWA was used to map reads against the reference assembly NCBI m37. Quality measures were calculated (Supplementary Excel File 8) as described for the human data with the exception that we did not require samples to have ≥ 1×10^6^ unique reads, because CKp25 samples were sequenced at a lower coverage. H3K9Ac domains were defined as regions that were covered by a peak in at least 4 (17%) out of the 23 samples (n=12 CKp25; n=11 tau) resulting in 44,165 domains. For each mouse model and time point, we compared 3 (or 2) AD model mice to 3 respective control mice using voom implemented in the R package limma^50^. Voom has been designed for small sample sizes and provides reliable domain-specific estimates of the variance by estimating the mean-variance trend using information from all domains. The voom method is incorporated in a linear regression framework with log_2_-transformed reads per million (log_2_ rpm) as outcome. A model was fitted for each domain and the cross correlation was added as technical covariate in addition the AD/control status variable.

### Induction of human neuronal cells overexpressing *MAPT*

The iPSC line used in this study (YZ1) was originally generated from the IMR-90 cell line (ATCC) and characterization of this line was described previously.^51^ Due to a karyotype abnormality in a small subset of cells, monoclonal isolates were obtained and confirmed to be karyotypically normal and pathogen free prior to initiating this study. The iPSC line used was confirmed to be of the correct identity prior to and at the conclusion of the study using short tandem repeat (STR) profiling (Genetica Cell Line Testing). Neurons were generated from the direct conversion of induced pluripotent stem cells by transduction with Neurogenin 2 as previously described^52^. Neurons were plated on Matrigel coated 96-well plates on DIV4 and maintained in media consisting of 485 mL neurobasal medium (Gibco), 5 mL Glutamax, 7.5 mL 20% Dextrose, 2.5 mL MEM NEAA with 1:50 B27, BDNF, CNTF, GDNF and doxcylcine. On DIV17, cells were transfected with a 1:1 dilution of lentivirus-packaged open reading frames (ORF) expressing human *MAPT* (titer 6.2×10^6^), resulting in overexpression of tau protein, or a GFP expressing construct (pRosetta). Following 18-hour incubation, virus-containing media was removed and replaced with fresh media to incubate until DIV31 when harvested for ATAC-seq. RNA-sequencing revealed a 10-fold increase of *MAPT* mRNA in cells transfected with *MAPT* cDNA compared to controls. In total, 9 *MAPT* OE and 9 control iNs were generated in 3 batches (differentiation rounds) of triplicate wells.

### 17-DMAG treatment of induced neurons

The second and third batch of experiments contained cell cultures of induced neurons treated with 1, 3, or 10 nM 17-DMAG for 24 hours prior to collection. Concentrations were chosen based on previous work (Gao et al., 2014). To account for vehicle specific effects, *MAPT* overexpressing cells and control cells cultured in concentration-matched DMSO solution for 24 hours prior to collection. Experimental design of the iN experiments is shown in Supplementary Table 5.

### Assay for transposase-accessible chromatin using sequencing (ATAC-seq)

ATAC-seq was performed as previously described^28^ with three modifications. (i) To reduce mitochondrial DNA contamination, we substituted 0.1% Ipegal with 0.01% Tween in the nuclear lysis buffer. (ii) Additional tagmentation buffer (TD; 2x) was prepared as follows: 20 mM Tris(hydroxymethyl)aminomethane and 10 mM MgCl2; adjusted to pH 7.6 with 100% acetic acid before addition of 20% (vol/vol) dimethylformamide^53^. (iii) We performed the PCR amplification in a 25ul total volume rather than 50ul to save half of our tagmented DNA for backup. After 31 days in vitro, iPSC neurons were trypsinised and counted to provide 50,000 live cells for ATAC-seq. iPSC-derived neurons were lysed with 0.01% Tween, 10 mM Tris-Cl, pH 7.4, 10 mM NaCl, 3 mM MgCl2 and nuclei pelleted, washed and transposed upon addition of 2.5 uL Nextera Tn5 Transposase in 1x TD buffer (Nextera Kit, Illumina) and incubation at 37°C for 30 mins. Immediately following transposition, DNA fragments were purified using a Qiagen MinElute PCR Purification Kit (Qiagen) following the manufacturer’s instructions. Transposed DNA was amplified with dual indexed Nextera PCR primers for 5 cycles prior to removing a 5 ul aliquot from each sample to test further amplification requirements by qPCR. Rounding up the cycle threshold from each sample decides how many additional cycles each sample needs to be amplified. For these samples, we needed to re-array and continue amplifying the remaining 20 ul for 5-8 cycles. Final libraries were cleaned twice with 1.5X Ampure XP SPRI beads and eluted in 12 ul of water. We quantified with Qubit HS DNA assay and assessed for quality using a Bioanalyzer High-Sensitivity DNA analysis Kit. 20 ng of each sample were pooled and diluted to 4nM using an estimated library length of 300 bp for all and submitted to the Broad Institute’s Genomics Platform for Illumina HiSeq2500 for 25 bp paired end reads.

### ATAC-seq data pre-processing and peak detection

Paired-end reads from ATAC-seq were aligned by the BWA algorithm against the human reference genome GRCh37^42^. MACS2 was applied without control library to each sample using fragment sizes obtained from the paired alignments and a q-value cutoff of 0.001^43^. Our ChIP-seq quality control pipeline was adapted for ATAC-seq: Instead of cross-correlation we calculated the median insert size and verified that the distribution of insert sizes showed a periodicity equal to the helical pitch of DNA (Supplementary Excel File 5). ATAC domains were defined as genomic regions covered by a peak in at least 4 (22%) of our 18 samples (n=9 *MAPT* OE; n=9 controls). Domains less than 50 bp away from each other were merged. In total, we obtained 40,637 ATAC domains. ATAC domains were annotated with chromatin states obtained from hESC-derived neurons included in the Roadmap Epigenomics Project (sample E010, core 15-state model)^11^. ATAC domains were clustered based on the proportion of chromatin states within the domains and classified into TSS, TSS flanking, enhancer or other domain as described in the Supplementary Notes (Supplementary Fig. 4a-c).

### Statistical analysis of ATAC-seq data

The number of read-pairs in each ATAC domain and sample were obtained and rpm values were calculated. The voom method was applied to test for different read counts in ATAC domains^50^. For each ATAC domain, log_2_-transformed rpm values were regressed on the tau status variable (n=9 *MAPT* OE, n=9 controls), the batch variable (3 batches with 6 samples each), and the logarithmized total number of reads obtained for the sample. The coefficient and p-value of the tau status variable was used to define genomic segments of consistent tau-related changes. To assess the reproducibility of the results between batches (Supplementary Fig. 4d-j), the same regression model but without the batch covariate was applied to each batch separately. This model was also used for analyzing the effect of 17-DMAG: Treated samples were compared to the respective control samples of the same batch for each 17-DMAG concentration separately while adjusting for different total numbers of reads. In this model, the coefficient for the tau status variable (n=3 *MAPT* OE + 17-DMAG, n=3 control) corresponds to the adjusted log_2_ fold change between the ATAC-seq reads in the 17-DMAG treated and the control samples. These coefficients were then used as outcome in a one-way ANOVA model with the 99 ATAC-seq derived segments as explanatory variable and the R^2^ value of this model was calculated to assess whether the location of an ATAC domain in a specific segment is associated with the magnitude of the tau effect.

### H3K9Ac Chip-seq data from induced neurons

H3K9Ac ChIP-seq data from iNs were generated and processed as described for the DLPFC tissue samples (Supplementary Excel File 5). Pooled input DNA from all three samples (n=2 *MAPT* OE, n=1 control) were used as control library during peak detection. Peaks that were detected in at least two of the three samples were defined as neuronal H3K9Ac domains (n=34,988) and used for subsequent analyses. The voom method was applied to test for differences between *MAPT* OE and control neurons^50^. In addition to the tau status variable (n=2 *MAPT* OE, n=1 control), cross-correlation was added as covariate to adjust for technical variability.

### Search for potential drugs in the Connectivity Map data base

Only H3K9Ac domains spanning a transcription start site of an active transcript (≥ 100 samples with rpkm ≥ 2 in the RNA-seq data) were considered. Gene annotation of the transcription start sites were used to map H3K9Ac domains to genes in the Connectivity Map data base^29^. The search algorithm provided by the Connectivity Map 2.0 requires a set of up- and downregulated genes as input. Following the manual, we selected the top 250 positively tau-associated genes and the top 250 negatively tau-associated genes that had the smallest p-values in our H3K9Ac data and calculated the enrichment score and p-values on the web server (https://www.broadinstitute.org/cmap). In addition, we calculated the Spearman rank correlation coefficient between the vector of standardized coefficients β _t_ /σ _βt_ for tau obtained from our H3K9Ac data and the differential expression statistic from the Connectivity Map for each compound (n=1309) in the data base. A negative correlation indicates that genes showing increased H3K9Ac levels with tau were repressed by the respective compound.

## Data availability

Human cortical ChIP-seq data was deposited at Synapse (Synape: syn4896408). Murine ChIP-seq data (GEO: GSE97560) and ATAC-seq data from iNeurons (GEO: GSE97409) are available at Gene Expression Omnibus.

## References

1. Lambert, J.C., et al. Meta-analysis of 74,046 individuals identifies 11 new susceptibility loci for Alzheimer’s disease. Nat Genet 45, 1452–1458 (2013).

2. Klein, H.U., Bennett, D.A. & De Jager, P.L. The epigenome in Alzheimer’s disease: current state and approaches for a new path to gene discovery and understanding disease mechanism. Acta neuropathologica 132, 503–514 (2016).

3. Lardenoije, R., et al. The epigenetics of aging and neurodegeneration. Progress in neurobiology 131, 21–64 (2015).

4. De Jager, P.L., et al. Alzheimer’s disease: early alterations in brain DNA methylation at ANK1, BIN1, RHBDF2 and other loci. Nat Neurosci 17, 1156–1163 (2014).

5. Lunnon, K., et al. Methylomic profiling implicates cortical deregulation of ANK1 in Alzheimer’s disease. Nat Neurosci 17, 1164–1170 (2014).

6. Frost, B., Bardai, F.H. & Feany, M.B. Lamin Dysfunction Mediates Neurodegeneration in Tauopathies. Current biology: CB 26, 129–136 (2016).

7. Frost, B., Hemberg, M., Lewis, J. & Feany, M.B. Tau promotes neurodegeneration through global chromatin relaxation. Nat Neurosci 17, 357–366 (2014).

8. Mansuroglu, Z., et al. Loss of Tau protein affects the structure, transcription and repair of neuronal pericentromeric heterochromatin. Scientific reports 6, 33047 (2016).

9. Bennett, D.A., Schneider, J.A., Arvanitakis, Z. & Wilson, R.S. Overview and findings from the religious orders study. Current Alzheimer research 9, 628–645 (2012).

10. Bennett, D.A., et al. Overview and findings from the rush Memory and Aging Project. Current Alzheimer research 9, 646–663 (2012).

11. Roadmap Epigenomics, C., et al. Integrative analysis of 111 reference human epigenomes. Nature 518, 317–330 (2015).

12. Reich, D.E., et al. Linkage disequilibrium in the human genome. Nature 411, 199–204 (2001).

13. Olshen, A.B., Venkatraman, E.S., Lucito, R. & Wigler, M. Circular binary segmentation for the analysis of array-based DNA copy number data. Biostatistics 5, 557–572 (2004).

14. Lieberman-Aiden, E., et al. Comprehensive mapping of long-range interactions reveals folding principles of the human genome. Science 326, 289–293 (2009).

15. Won, H., et al. Chromosome conformation elucidates regulatory relationships in developing human brain. Nature 538, 523–527 (2016).

16. Meuleman, W., et al. Constitutive nuclear lamina-genome interactions are highly conserved and associated with A/T-rich sequence. Genome research 23, 270–280 (2013).

17. Kind, J., et al. Genome-wide maps of nuclear lamina interactions in single human cells. Cell 163, 134–147 (2015).

18. Klein, H.U. & De Jager, P.L. Uncovering the Role of the Methylome in Dementia and Neurodegeneration. Trends in molecular medicine 22, 687–700 (2016).

19. Schoofs, T., et al. DNA methylation changes are a late event in acute promyelocytic leukemia and coincide with loss of transcription factor binding. Blood 121, 178–187 (2013).

20. Zou, F., et al. Brain expression genome-wide association study (eGWAS) identifies human disease-associated variants. PLoS Genet 8, e1002707 (2012).

21. Houseman, E.A., et al. DNA methylation arrays as surrogate measures of cell mixture distribution. BMC bioinformatics 13, 86 (2012).

22. Jaffe, A.E. & Irizarry, R.A. Accounting for cellular heterogeneity is critical in epigenome-wide association studies. Genome Biol 15, R31 (2014).

23. Liang, W.S., et al. Alzheimer’s disease is associated with reduced expression of energy metabolism genes in posterior cingulate neurons. Proceedings of the National Academy of Sciences of the United States of America 105, 4441–4446 (2008).

24. Yoshiyama, Y., et al. Synapse loss and microglial activation precede tangles in a P301S tauopathy mouse model. Neuron 53, 337–351 (2007).

25. Cruz, J.C., Tseng, H.C., Goldman, J.A., Shih, H. & Tsai, L.H. Aberrant Cdk5 activation by p25 triggers pathological events leading to neurodegeneration and neurofibrillary tangles. Neuron 40, 471–483 (2003).

26. Gjoneska, E., et al. Conserved epigenomic signals in mice and humans reveal immune basis of Alzheimer’s disease. Nature 518, 365–369 (2015).

27. Muratore, C.R., et al. Cell-type dependent Alzheimer’s disease phenotypes: probing the biology of selective neuronal vulnerability. Stem Cell Reports (2017).

28. Buenrostro, J.D., Giresi, P.G., Zaba, L.C., Chang, H.Y. & Greenleaf, W.J. Transposition of native chromatin for fast and sensitive epigenomic profiling of open chromatin, DNA-binding proteins and nucleosome position. Nat Methods 10, 1213–1218 (2013).

29. Lamb, J., et al. The Connectivity Map: using gene-expression signatures to connect small molecules, genes, and disease. Science 313, 1929–1935 (2006).

30. Dickey, C.A., et al. The high-affinity HSP90-CHIP complex recognizes and selectively degrades phosphorylated tau client proteins. J Clin Invest 117, 648–658 (2007).

31. Luo, W., et al. Roles of heat-shock protein 90 in maintaining and facilitating the neurodegenerative phenotype in tauopathies. Proceedings of the National Academy of Sciences of the United States of America 104, 9511–9516 (2007).

32. Gao, L., et al. Discovery of the neuroprotective effects of alvespimycin by computational prioritization of potential anti-Parkinson agents. The FEBS journal 281, 1110–1122 (2014).

33. Imakaev, M., et al. Iterative correction of Hi-C data reveals hallmarks of chromosome organization. Nat Methods 9, 999–1003 (2012).

34. Dixon, J.R., et al. Chromatin architecture reorganization during stem cell differentiation. Nature 518, 331–336 (2015).

35. Fortin, J.P. & Hansen, K.D. Reconstructing A/B compartments as revealed by Hi-C using long-range correlations in epigenetic data. Genome Biol 16, 180 (2015).

36. Nora, E.P., et al. Spatial partitioning of the regulatory landscape of the X-inactivation centre. Nature 485, 381–385 (2012).

37. Dixon, J.R., et al. Topological domains in mammalian genomes identified by analysis of chromatin interactions. Nature 485, 376–380 (2012).

38. Ong, C.T. & Corces, V.G. CTCF: an architectural protein bridging genome topology and function. Nature reviews. Genetics 15, 234–246 (2014).

39. Chang, K.H., et al. Nuclear envelope dispersion triggered by deregulated Cdk5 precedes neuronal death. Molecular biology of the cell 22, 1452–1462 (2011).

40. Bennett, D.A., Schneider, J.A., Tang, Y., Arnold, S.E. & Wilson, R.S. The effect of social networks on the relation between Alzheimer’s disease pathology and level of cognitive function in old people: a longitudinal cohort study. The Lancet. Neurology 5, 406–412 (2006).

41. Bennett, D.A., Schneider, J.A., Wilson, R.S., Bienias, J.L. & Arnold, S.E. Neurofibrillary tangles mediate the association of amyloid load with clinical Alzheimer disease and level of cognitive function. Archives of neurology 61, 378–384 (2004).

42. Li, H. & Durbin, R. Fast and accurate short read alignment with Burrows-Wheeler transform. Bioinformatics 25, 1754–1760 (2009).

43. Zhang, Y., et al. Model-based analysis of ChIP-Seq (MACS). Genome Biol 9, R137 (2008).

44. Landt, S.G., et al. ChIP-seq guidelines and practices of the ENCODE and modENCODE consortia. Genome research 22, 1813–1831 (2012).

45. Lun, A.T. & Smyth, G.K. De novo detection of differentially bound regions for ChIP-seq data using peaks and windows: controlling error rates correctly. Nucleic acids research 42, e95 (2014).

46. Lim, A.S., et al. 24-hour rhythms of DNA methylation and their relation with rhythms of RNA expression in the human dorsolateral prefrontal cortex. PLoS Genet 10, e1004792 (2014).

47. Li, B. & Dewey, C.N. RSEM: accurate transcript quantification from RNA-Seq data with or without a reference genome. BMC bioinformatics 12, 323 (2011).

48. Hebestreit, K., Dugas, M. & Klein, H.U. Detection of significantly differentially methylated regions in targeted bisulfite sequencing data. Bioinformatics 29, 1647–1653 (2013).

49. Allen, M., et al. Human whole genome genotype and transcriptome data for Alzheimer’s and other neurodegenerative diseases. Scientific data 3, 160089 (2016).

50. Law, C.W., Chen, Y., Shi, W. & Smyth, G.K. voom: Precision weights unlock linear model analysis tools for RNA-seq read counts. Genome Biol 15, R29 (2014).

51. Zeng, H., et al. Specification of region-specific neurons including forebrain glutamatergic neurons from human induced pluripotent stem cells. PLoS One 5, e11853 (2010).

52. Zhang, Y., et al. Rapid single-step induction of functional neurons from human pluripotent stem cells. Neuron 78, 785–798 (2013).

53. Wang, Q., et al. Tagmentation-based whole-genome bisulfite sequencing. Nature protocols 8, 2022–2032 (2013).

